# *Toxoplasma gondii* co-opts the unfolded protein response to enhance migration and dissemination of infected host cells

**DOI:** 10.1101/2020.04.14.042069

**Authors:** Leonardo Augusto, Jennifer Martynowicz, Parth H. Amin, Nada S. Alakhras, Mark H. Kaplan, Ronald C. Wek, William J. Sullivan

**Author notes:** Corresponding authors: Ronald C. Wek, Showalter Professor, Indiana University School of Medicine, 635 Barnhill Drive, MS 4067A, Indianapolis, IN 46202, William J. Sullivan, Jr., Showalter Professor, Indiana University School of Medicine, 635 Barnhill Drive, MS A418C, Indianapolis, IN 46202.

## Abstract

*Toxoplasma gondii* is an intracellular parasite that reconfigures its host cell to promote pathogenesis. One consequence of *Toxoplasma* parasitism is increased migratory activity of host cells, which facilitates dissemination. Here we show that *Toxoplasma* triggers the unfolded protein response (UPR) in host cells through calcium release from the endoplasmic reticulum (ER). We further found that host IRE1, an ER stress sensor protein activated during *Toxoplasma* infection, also plays a noncanonical role in actin remodeling by binding filamin A in infected cells. By inducing cytoskeletal remodeling via IRE1 oligomerization in host cells, *Toxoplasma* enhances host cell migration *in vitro* and dissemination of the parasite to host organs *in vivo*. Our study identifies novel mechanisms used by *Toxoplasma* to induce dissemination of infected cells, providing new insights into strategies for treatment of toxoplasmosis.

**Importance:** Cells that are infected with the parasite *Toxoplasma gondii* exhibit heightened migratory activity, which facilitates dissemination of the infection throughout the body. In this study, we identify a new mechanism used by *Toxoplasma* to hijack its host cell and increase its mobility. We further show that the ability of *Toxoplasma* to increase host cell migration does not involve the enzymatic activity of IRE1, but rather IRE1 engagement with actin cytoskeletal remodeling. Depletion of IRE1 from infected host cells reduces their migration in vitro and significantly hinders dissemination of *Toxoplasma* in vivo. Our findings reveal a new mechanism underlying host-pathogen interactions, demonstrating how host cells are co-opted to spread a persistent infection around the body.

## Introduction

*Toxoplasma gondii* is an obligate intracellular parasite capable of infecting any nucleated cell in warm-blooded vertebrates. Recent studies have revealed a striking degree of host cell remodeling taking place in *Toxoplasma*-infected cells that serves to facilitate pathogenesis and transmission. In addition to secreted parasite effectors that modulate host cell gene expression, *Toxoplasma* infection can alter immune responses and enable dissemination to other host tissues [1]. Therein, *Toxoplasma* can differentiate from the replicative tachyzoites to the latent bradyzoite stage, enabling formation of tissue cysts that persist for the lifetime of the infected host [2].

Upon host cell invasion, *Toxoplasma* forms a parasitophorous vacuole (PV) that serves as a protective niche that can interface with the host cell cytoplasm to sequester nutrients [3]. Curiously, *Toxoplasma* recruits the host endoplasmic reticulum (ER) to the PV via association between their respective membranes, although the reasons for this high affinity interaction are not yet understood [4, 5].

The ER is sensitive to the perturbations in protein homeostasis through a stress-sensing pathway known as the unfolded protein response (UPR). Three ER transmembrane proteins, IRE1, ATF6, and PERK operate as sensors that activate the UPR, leading to changes in gene expression that restore and expand the processing capacity of the organelle [6-8]. IRE1 (ERN1) is a protein kinase and endoribonuclease that facilitates cytosolic splicing of *XBP1* (XBP1s) mRNA, thereby enhancing expression of the XBP1s isoform, which induces transcription of genes involved in ER-associated protein degradation (ERAD), lipid synthesis, and protein folding [7, 8]. In response to ER stress, ATF6 transits from the ER to the Golgi apparatus where it is cleaved, releasing an N-terminal cytosolic fragment (ATF6-N) that enters the nucleus and activates UPR-target genes involved in protein folding and transport [6, 9]. PERK (EIF2AK3) is the third UPR sensor, which phosphorylates eIF2α to direct translational and transcriptional modes of gene expression that regulate ER processing of proteins, metabolism, and the oxidation status of cells [6, 10]. While the three ER stress sensory proteins function in parallel, there is cross-regulation that serves to coordinate the timing and magnitude of the UPR. For example, PERK was reported to induce expression of RPAP2, which serves to dephosphorylate and repress IRE1, thereby providing a means for the cell to abort failed ER-stress adaptation and trigger apoptosis [11].

In addition to its role in the UPR, IRE1 was recently shown to modulate cytoskeletal remodeling and cell migration through direct interactions with the actin crosslinking factor filamin A [12]. The role of IRE1 in cytoskeletal remodeling is enhanced by pharmacological induction of ER stress, but occurs independent of IRE1 protein kinase and endoribonuclease activities [12]; rather, IRE1 serves as a scaffolding protein for filamin A to orchestrate changes in cellular motility. This is noteworthy because *Toxoplasma* stimulates host cell migration, turning its host cell into a “Trojan Horse” that can ferry parasites throughout the body [13]. Given the recruitment of host ER to the PV, we postulated that migratory activities mediated by IRE1 function in parasite dissemination. In the present study, we uncover a new mechanism by which *Toxoplasma* alters host ER homeostasis to produce hypermigratory activity in infected host cells. We show that during the course of *Toxoplasma* infection, the three UPR sensory proteins in the host cells are activated by a process involving calcium release from the ER, leading to IRE1 oligomerization, association with filamin A, and enhanced cell migration. Importantly, the IRE1-associated migration is a crucial determinant for successful dissemination of toxoplasmosis in a mouse model of infection.

## Results

### Induction of the UPR in *Toxoplasma*-infected host cells

It is currently unclear why intracellular tachyzoites recruit host ER to the parasite PV. To address whether *Toxoplasma* perturbs host ER homeostasis, we infected mouse embryonic fibroblast (MEF) cells with RH strain parasites and measured three primary markers of the host UPR over a 36-hour time course. Within 12 h of infection, *Toxoplasma* increased activation of PERK as measured by its self-phosphorylation (PERK-P), induced expression of ATF6 and formation of its cleavage product ATF6-N, and increased levels of the IRE1-derived spliced variant of XBP1 (XBP1s) **(Fig. 1A)**. It is noteworthy that whereas expression of ATF6-N and XBP1s were transient, with increased amounts of the proteins appearing between 12 to 20 h post-infection (hpi), PERK-P increased throughout the 36 h of infection **(Fig. 1A)**. These results indicate that *Toxoplasma* infection causes ER stress that activates each of the sensory proteins of the UPR with some differences in the duration of their induction.

**Figure. 1.**
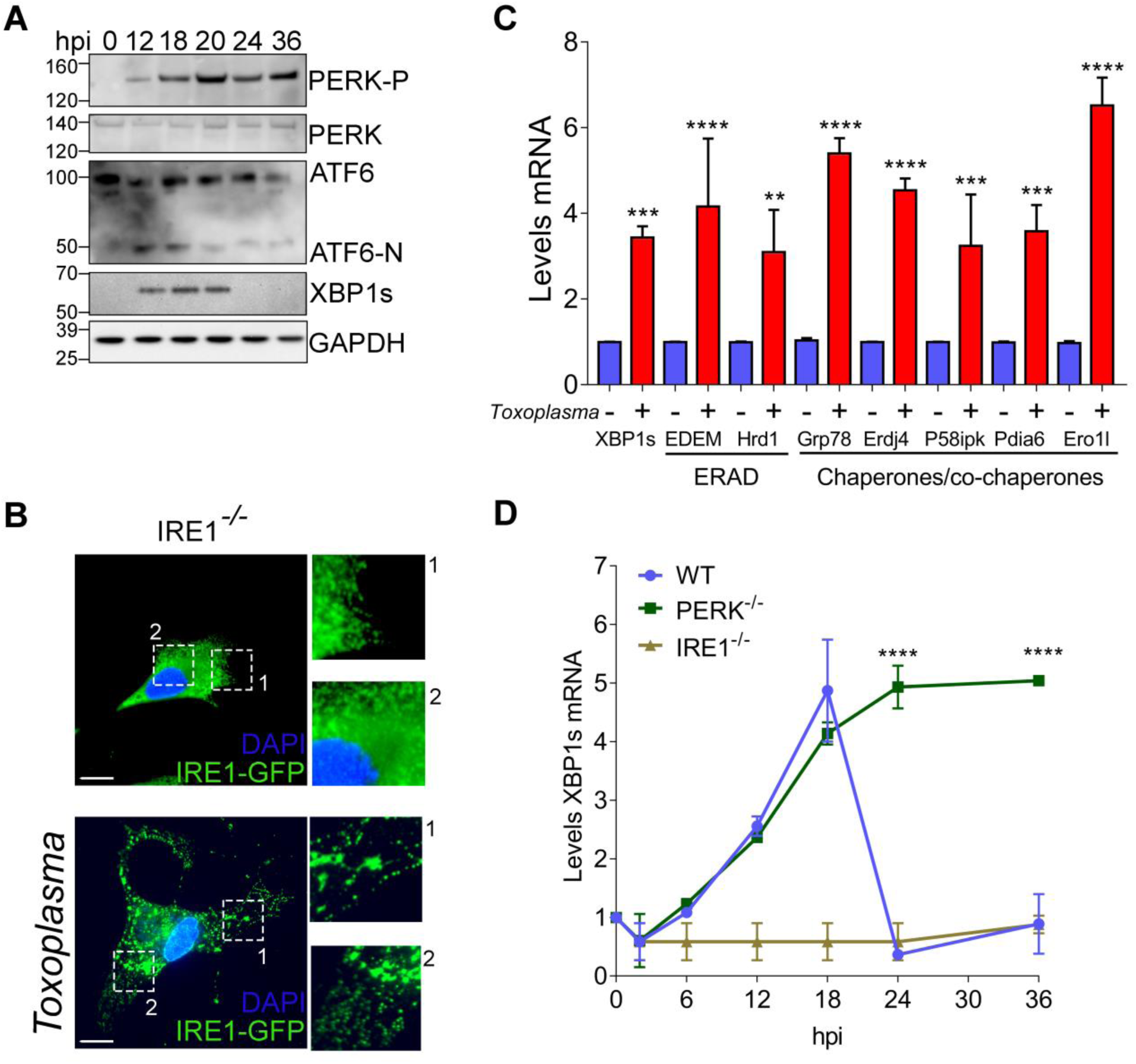
*Toxoplasma* infection triggers activation of the UPR in host cells. (A) At the indicated times following infection with *Toxoplasma*, cells were harvested and the levels of total PERK, PERK-P, full length ATF6 and ATF6-N, XBP1s, and GAPDH were measured by immunoblot analyses. (B) *IRE1*^-/-^ MEF cells were transfected with a plasmid encoding EGFP- IRE1 (green), followed by infection with *Toxoplasma*. DAPI (blue) was used to visualize host cell and parasites nuclei. Two boxed areas (1 and 2) of each condition are amplified to highlight the IRE1 distribution. Bar=5 µm. (C) MEF cells were infected with *Toxoplasma* for 18 h, or were mock infected, and the mRNA levels of the indicated ERAD/chaperone genes were measured by RT-qPCR. Levels of mRNA were normalized to mock infected, which is represented as a value of 1. (±SD, n=3) ***p<0.001, ****p< 0.0001. (D) *XBP1s* mRNA levels were measured by RT-qPCR at the indicated hpi in WT, *PERK*^-/-^ and *IRE1*^-/-^ MEF cells, as indicated. Levels of *XBP1s* mRNA were normalized to total *XBP1* mRNA in mock infected cells (value of 1) at each time point. (±SD, n=3) ****p<0.0001.

Activation of IRE1 involves oligomerization that can be visualized by a pattern of punctate spots by IFA [14-16]. We expressed EGFP-tagged IRE1 in MEF cells that were deleted for the endogenous *IRE1* gene by CRISPR/Cas9 genome editing **(Fig. S1A-C)**; upon *Toxoplasma* infection, we observed formation of IRE1 foci that is consistent with reported IRE1 activation by oligomerization **(Fig. 1B)** [15, 16]. Furthermore, expression of *XBP1s* mRNA, and its downstream target genes involved in ERAD and protein folding, were induced upon *Toxoplasma* infection of wild-type (WT) MEF cells **(Fig. 1C)**. These results indicate that IRE1 activation and signaling occur in response to *Toxoplasma* infection.

To further study the timing of UPR induction and potential cross-regulation between the UPR sensors in infected cells, we measured *XBP1s* mRNA levels by RT-qPCR in WT, *IRE1*^*-/-*^, or *PERK*^*-/-*^ MEF cells. In WT cells, levels of *XBP1s* mRNA rose sharply until 18 hpi **(Fig. 1D)**, consistent with the increase in XBP1s protein **(Fig. 1A)**. As expected, *XBP1s* mRNA was not detected in infected *IRE1*^-/-^ cells **(Fig. 1D)**. By comparison, there was induced *XBP1s* mRNA that was sustained during 36 h of *Toxoplasma* infection in *PERK*^*-/-*^ cells, indicating that without PERK, IRE1 continues to facilitate *XBP1s* expression **(Fig. 1D)**. These results are consistent with the idea that PERK governs IRE1 activity as previously reported [11] and suggests that in infected WT cells, PERK operates to attenuate the response of IRE1 after 18 hpi.

### IRE1 affects calcium release from ER in *Toxoplasma*-infected cells

Protein folding in the ER is highly sensitive to the concentration of calcium, which is released from the organelle by ryanodine receptors (RyR) and inositol 1,4,5-triphosphate (IP3)- receptors (IP_3_R) [17]. The ER is a major reservoir of calcium; disruptions of calcium homeostasis can lead to unfolded proteins and initiation of the UPR [17]. To determine whether calcium content is altered in the host ER during *Toxoplasma* infection, we first measured cytosolic calcium in infected MEF cells. Over an 18-hour period, *Toxoplasma* infection induced a steady increase in host cell cytosolic calcium levels **(Fig. 2A, Fig. S2A)**. Basal calcium levels in the cytosol of *IRE1*^-/-^ cells were lower compared to WT cells, as previously reported [18], with some increase upon parasite infection **(Fig. 2A, Fig. S2A)**. To assess the mode of calcium release from the host ER during infection, we monitored calcium transport using Fluo-4AM in the presence of antagonists of RyR or IP_3_R. The cytosolic calcium levels were lower in infected cells treated with either antagonist, with the most robust calcium reduction occurring with inhibition of IP_3_R **(Fig. 2B)**. We further addressed the contribution of RyR and IP_3_R to calcium release by adding increasing doses of caffeine and IP_3_, agonists of RyR and IP_3_R receptor activity [19], respectively, to WT and *IRE1*^-/-^ cells infected with *Toxoplasma* for 18 h and incubated with Mag-Fluo-4. To compare calcium release between RyR and IP_3_R in WT and *IRE1*^-/-^ cells, the fluorescence values were represented as the percentage of calcium release and the respective start points were normalized to untreated WT and *IRE1*^-/-^ cells, respectively. The RyR agonist enhanced calcium to similar levels in infected WT or *IRE1*^-/-^ cells **(Fig. 2C)**. By comparison, the agonist IP_3_ induced appreciable calcium release only in infected WT cells **(Fig. 2D)**, indicating a role for IRE1 in regulating IP_3_R activity as previous reported [18].

**Figure. 2.**
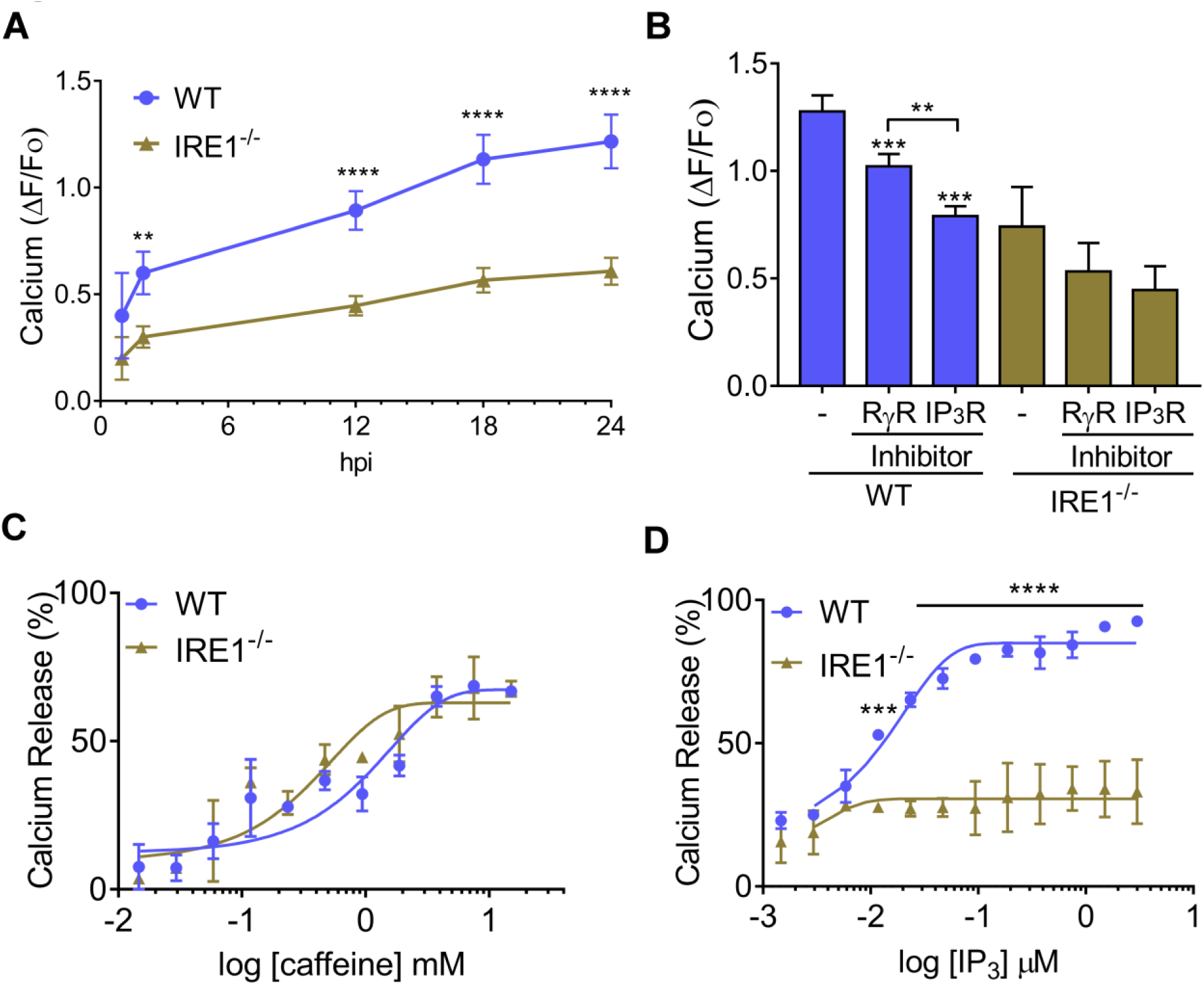
*Toxoplasma* infection induces calcium release. (A) At the indicated time hpi, cytosolic calcium levels were measured in the infected WT and *IRE1*^-/-^ cells. Values of infected cells were normalized to mock-infected cells (±SD, n=3), **p<0.005 and ****p<0.001. (B) Infected WT and *IRE1*^-/-^ cells were treated with RyR or IP_3_R inhibitors for 6 h and the levels of cytosolic calcium were measured. 100 µM Ryanodine (RyR inhibitor) and 0.6 µM Xestospongin C (XeC) (IP_3_R inhibitor), (±SD, n=3) **p<0.05, ***p<0.005. Infected cells were incubated with Mag-Fluo-4, followed by plasma membrane permeabilization with saponin and incubation with ATP to maintain the calcium in the ER. To estimate RyR (C) and IP_3_R (D) activity, WT and *IRE1*^-/-^ cells infected with *Toxoplasma* for 18 h were treated with caffeine or IP_3_ at the indicated concentrations and calcium release is represented the fluorescence using Mag-Fluo-4 as described [19]. Values are represented as the percentage of calcium release and the respective start points were normalized to untreated WT and *IRE1*^-/-^ cells, respectively. (±SD, n=3), **p<0.05, ****p<0.001.

Surprisingly, the percentage of calcium release was higher when IP_3_R was stimulated compared to RyR, demonstrating that *Toxoplasma* infection differentially alters RyR and IP_3_R activities. Collectively, these results suggest that *Toxoplasma* infection induces significant calcium release from the host ER by processes involving both IP_3_R and RyR; this calcium release is influenced by IRE1 and is a likely contributor to the activation of the host ER-stress sensor proteins seen in Figure 1A.

### IRE1 activation induces cell migration in infected cells

*Toxoplasma* triggers rapid morphological changes in host cells, including disappearance of podosome structures and appearance of lamellipodia [20]. IRE1 has recently been shown to have noncanonical functions in actin cytoskeletal remodeling by directly binding to filamin A [12]. To address whether activation of IRE1 by *Toxoplasma* infection enhances host cell migration, we quantified the number of lamellipodia per infected cell normalized to uninfected cells **(Fig. 3A, Fig. S3A)**. At 18 hpi, *Toxoplasma* infection increased the number of lamellipodia in WT cells and these structures were significantly diminished in IRE1-deficient cells **(Fig. 3A)**. By comparison, there were greater numbers of lamellipodia in PERK-deficient cells or those treated with a PERK inhibitor, consistent with the idea that PERK is a negative regulator of IRE1 **(Fig. 3A)**. Of interest, treatment with inhibitors of IRE1, namely 4µ8c, which interferes with endoribonuclease activity, and KIRA6, which blocks IRE1 protein kinase activity, did not change the number of lamellipodia compared infected cells treated with vehicle **(Fig. 3A)**. These results indicate that IRE1 can control migration of *Toxoplasma* infected cells independent of its known enzymatic activities.

**Figure. 3.**
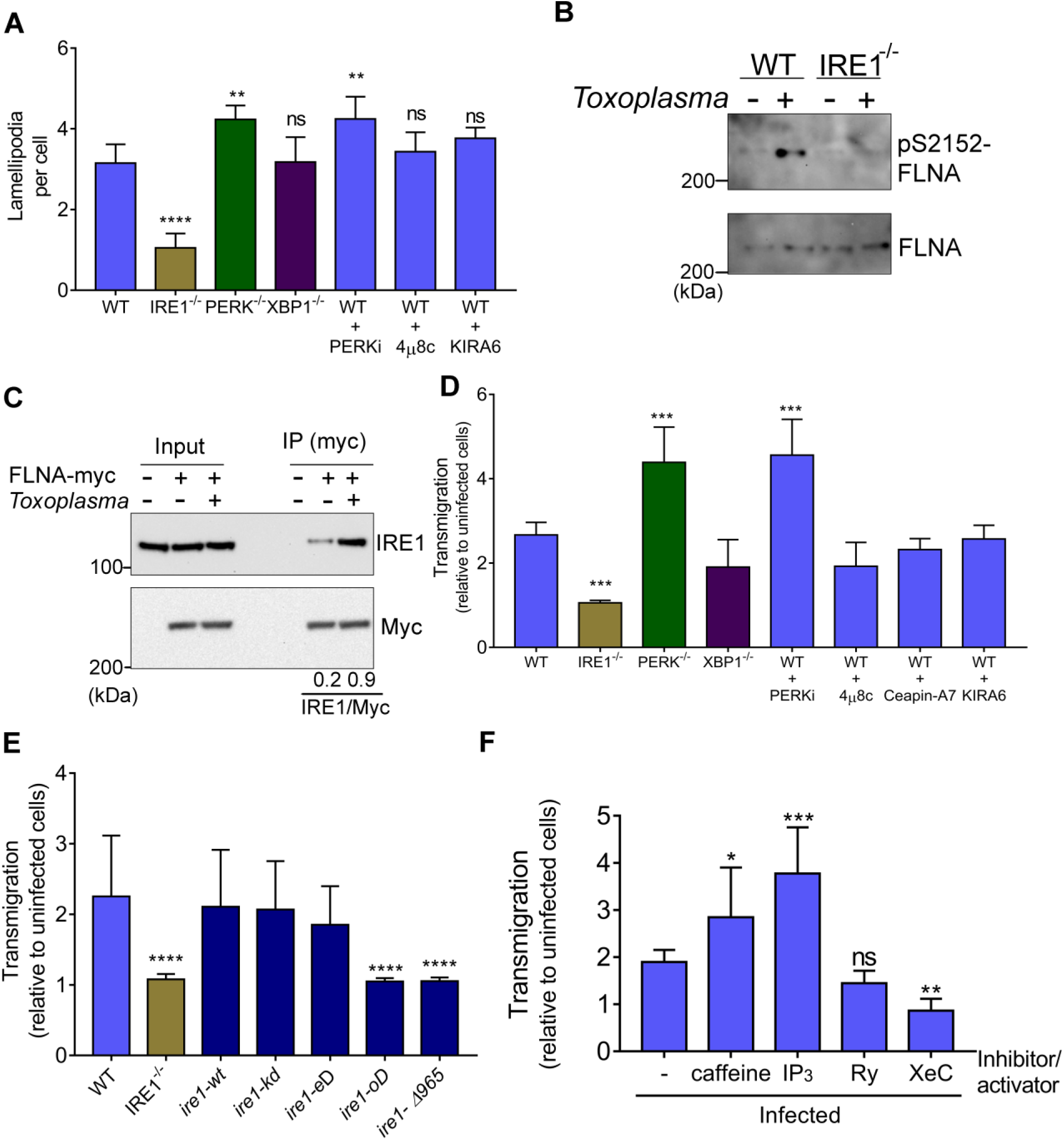
Activation of IRE1 enhances migration of cells infected with *Toxoplasma*. (A) The numbers of lamellipodia were determined in 50 randomly selected MEF cells infected with *Toxoplasma* 18 hpi and normalized to uninfected cells. Also shown are infected WT cells treated with 1.2 µM PERKi (GSK2656157), 0. 4 µM 4µ8c, 250 nM KIRA6, or 0.2 µM ceapin-A7 for 18 h. (±SD, n=5) **p<0.05, ***p<0.005. (B) WT and *IRE1*^*-/-*^ cells were infected with *Toxoplasma* for 18 h, then cells were harvested and the levels of filamin A phosphorylation (S2152) were measured by immunoblot analyses. (C) WT cells transiently expressing Myc-filamin A were infected with *Toxoplasma* for 18 h. IP of the tagged Filamin A was carried out using Myc magnetic beads. Bound proteins were separated by SDS-PAGE and the levels of Myc-filamin A and associated IRE1 were measured by immunoblot and compared to uninfected cells. Densitometry of IRE1 signal divided by Myc signal (IRE1/Myc). (D) WT or *IRE1*^*-/-*^, *PERK*^*-/-*^, and *XBP1*^*-/-*^ cells were infected with *Toxoplasma* for 18 h. Alternatively, WT cells were infected with parasite and treated with the following inhibitors during migration per 18 h: 1.2 µM PERKi (GSK2656157), 0.2 µM Ceapin-A7, 250 nM KIRA6, or 0.4 µM 4µ8c. Infected and mock infected cells were trypsinized, counted, and the same number of cells was used for the transmigration assay. Transmigration was determined by counting the number of infected normalized to noninfected cells that migrated through the membrane. (±SD n=3), *** p<0.0005. (E) *IRE1*^-/-^ cells were rescued with *ire1-wt* (wild-type), *kD* (kinase domain-dead- K599A), *eD* (endoribonuclease domain-dead- P830L), *oD* (oligomerization domain-dead- D123P) or *Δ965* (truncated c-terminal *Δ965*), and then a transmigration assay was carried out as described above. (±SD, n=3), ****p<0.0001. (F) Transmigration assay of infected WT MEF cells was carried out in the presence of IP_3_R and RyR inhibitors (0.6 µM Xestospongin C, XeC or 100 µM Ryanodine, Ry) or activators (100 µM IP_3_ or 1 mM caffeine). (±SD, n=3), *p<0.1, **p<0.05 and ***p<0.0005. ns = not significant.

The functions of filamin A in cytoskeleton dynamics and cell migration is dependent on phosphorylation of serine 2,152 (S2152) [21]. We detected a sharp increase in filamin A phosphorylation in MEF cells infected with *Toxoplasma* for 18 h, which was not observed in host cells lacking IRE1 **(Fig. 3B)**. To address whether there is an association between IRE1 and filamin A, we expressed myc-tagged filamin A (myc-FLNA) in WT MEF cells. Following 18 h of *Toxoplasma* infection, we then performed an immunoprecipitation (IP) of the tagged filamin A, followed by immunoblot measurements of associated IRE1. There was enhanced association of IRE1 with filamin A in cells infected with the parasite compared to those uninfected **(Fig**. **3C)**.

Next, we measured changes in host cell transmigration upon parasite infection and determined that there was a 2-fold increase in migration of WT MEF cells upon infection with *Toxoplasma* **(Fig. 3D, Fig. S3B)**. Increased host cell migration was observed regardless of whether the cells were infected with type I RH or type II ME49 strain parasites **(Fig. S3C)**. In contrast, parasite infection did not induce migration of cells lacking IRE1. Levels of migration in infected WT MEF cells treated with the IRE1 enzymatic inhibitors, KIRA6 or 4µ8c, or an ATF6 inhibitor (Ceapin-A7), were not significantly changed compared to untreated infected MEF cells, nor were they altered in cells lacking the downstream IRE1 target XBP1 **(Fig. 3D)**. Notably, *Toxoplasma*-induced migration of PERK-deficient cells or WT treated with PERK inhibitor was increased >2.5-fold upon parasite infection compared to infected cells with functional PERK **(Fig. 3D)**. These results suggest that IRE1 plays a critical role in inducing migration of *Toxoplasma*-infected cells and that this migration is independent of the protein kinase and endoribonuclease activities of IRE1 and its downstream target *XBP1*. Furthermore, PERK is suggested to dampen both IRE1 functions in *XBP1* mRNA splicing and cell migration, which are induced upon *Toxoplasma* infection.

### ER stress induces cell migration by mechanisms involving IRE1 oligomerization

To better understand the mechanisms by which IRE1 enhances host cell migration in response to *Toxoplasma* infection, we rescued the *IRE1*^-/-^ cells by expressing WT or defined mutant versions of IRE1 **(Fig. 3E)**. Equal amounts of the IRE1 proteins were expressed as judged by immunoblot and immunofluorescence analyses **(Fig. S4A and B)**. As expected, *ire1- wt* rescued the migration capacity of infected cells **(Fig. 3E)**. Expression of IRE1 defective in kinase (*ire1-kD*) or endoribonuclease (*ire1-eD*) activities still rescued the migration phenotype, further supporting the idea that these activities are dispensable for induced cell migration in response to parasite infection **(Fig. 3E)**. In contrast, cells expressing IRE1 with mutations in the oligomerization domain (*ire1-oD*) were deficient in *Toxoplasma*-induced migration **(Fig. 3E)**. Furthermore, a truncated c-terminal version of IRE1 (*ire1-Δ965)* lacking the proline-rich carboxy-terminal segment of IRE1 that binds filamin A [12] was also impaired in cell migration following infection **(Fig. 3E)**. As anticipated, only *ire1-wt* and *ire1-Δ965* showed induction of spliced *XBP1* mRNA upon pharmacological induction of ER stress **(Fig. S4C)**. These results suggest that *Toxoplasma* infection induces IRE1 oligomerization (see also Fig. 1B), and that the hypermigratory behavior of infected host cells is reliant on both the oligomerization domain and the portion of IRE1 that interacts with filamin A.

Since we found that *Toxoplasma* induces calcium release from the host ER upon infection, we examined the importance of calcium release in host cell migration induced by *Toxoplasma*. We treated infected cells with RyR and IP_3_R receptor blockers or activators during the migration assay: ryanodine (Ry) or xestospongin-C (XeC) were used to inhibit RyR and IP_3_R receptors, respectively; caffeine or IP_3_ were used to activate RyR and IP_3_R receptors, respectively. When infected cells were treated with the RyR and IP_3_R activators (releasing calcium from ER into the cytosol), the migration levels increased compared to cells not treated with these agents. By contrast, addition of the IP_3_R inhibitor significantly decreased migration of the infected cells **(Fig. 3F)**, consistent with IP_3_R being the major calcium release receptor involved in triggering the host UPR following *Toxoplasma* infection **(Fig. 2B)**. These results support the idea that calcium release from the host ER contributes to IRE1 activation and its subsequent role in augmenting migration in response to infection.

### IRE1 controls host cell migration in infected immune cells *in vitro*

*Toxoplasma* makes use of immune cells as a “Trojan Horse” to disseminate to distal organs and tissues throughout the body of the infected host [13]. To address whether *Toxoplasma* is targeting IRE1 in immune cells to enhance their migration and facilitate dissemination, we infected bone marrow-derived dendritic cells (DCs) with *Toxoplasma*. Levels of *XBP1s* mRNA were sharply increased upon *Toxoplasma* infection in DCs **(Fig. 4A)**, consistent with the idea that the parasite infection activated IRE1 in this cell type. Next, we used CRISPR/Cas9 and two distinct sgRNAs (sgRNA 1 and 2) to disrupt IRE1 in DCs **(Fig. S5A)**. The sgRNA1 and sgRNA2 decreased *IRE1* mRNA levels by 66% and 93%, respectively **(Fig. 4B)**. Each gRNA also led to a corresponding reduction in IRE1 protein in DCs, with sgRNA2 leading to no detectable IRE1 **(Fig. 4C)**. IRE1-depleted DCs (ire1 (-)) did not exhibit decreased cell viability compared to WT, nor did they show any difference in infectivity with *Toxoplasma* **(Fig. 4D, Fig. S5B)**. However, loss of IRE1 in DCs significantly lowered the transmigratory capacity following infection with either Type I or Type II strains of *Toxoplasma* **(Fig. 4E, Fig. S3D)**. To determine whether IP_3_R plays a role in the migration of infected DCs, we carried out the migration assay in the presence of XeC. Inhibition of IP_3_R resulted in a loss of host cell migration following 18 h of *Toxoplasma* infection **(Fig. 4F)**, further supporting the importance of calcium homeostasis in the ER for migration of *Toxoplasma* infected cells. To test whether IRE1 controls the migration of infected macrophages as well, we used CRISPR/Cas9 to deplete IRE1 in J774.1 macrophages **(Fig. 4G and H)**. As observed for DCs, the loss of IRE1 significantly reduced the ability of infected macrophages to migrate **(Fig. 4I)**.

**Figure. 4.**
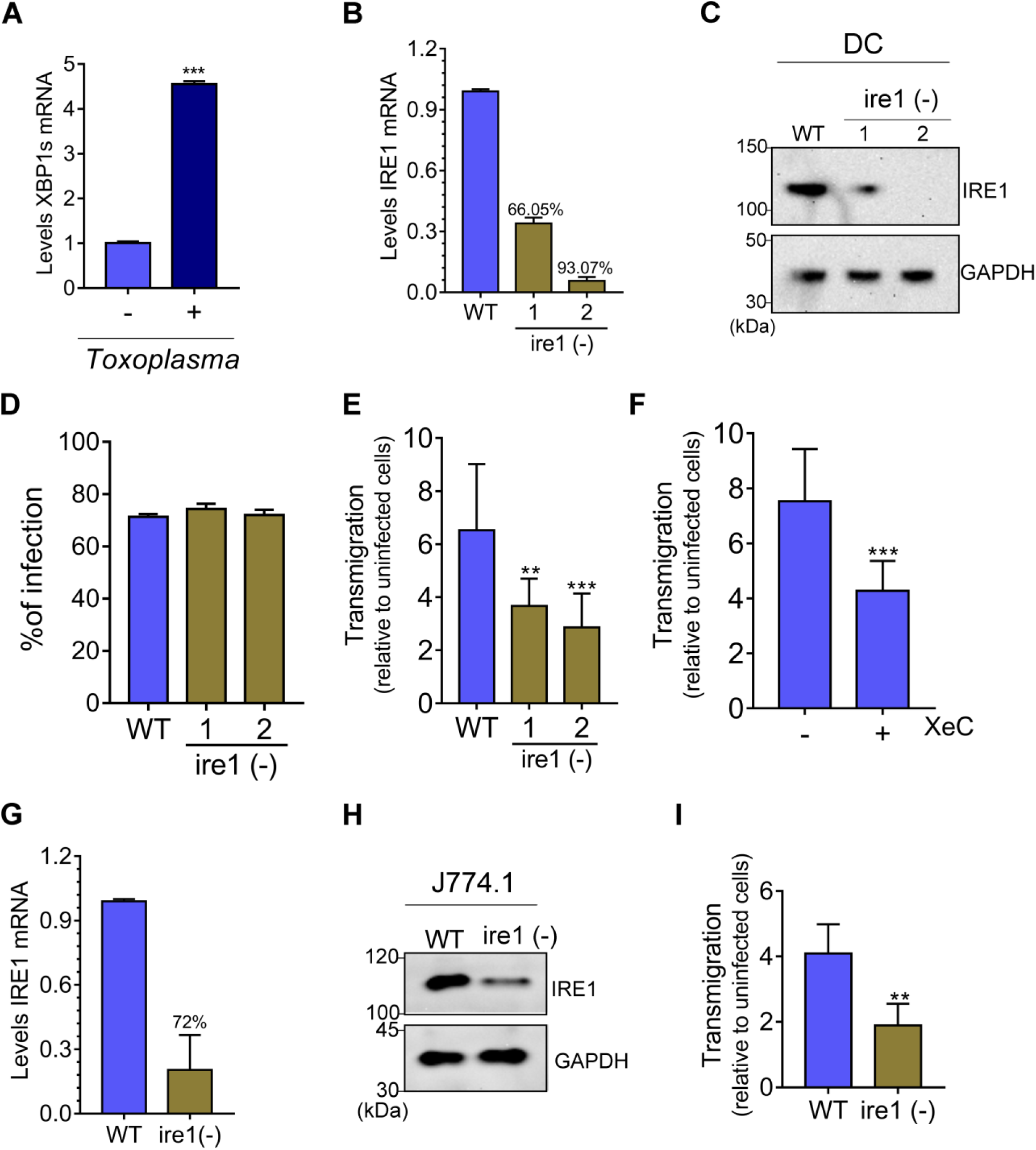
IRE1 is important for migration of infected DCs. (A) Bone marrow-derived DCs were infected with *Toxoplasma* for 18 h and *XBP1s* mRNA levels were measured by RT-qPCR. The values of XBP1s were normalized to values of total XBP1. (±SD, n=3) ***p<0.0005. (B-C) The CRISPR/Cas9-engineered depletion of IRE1 in DCs, designated ire (-), was assayed by RT- qPCR and immunoblot using IRE1 antibody compared to WT cells (GAPDH was included as a loading control). (D) Percentage of infection was determined by counting the number of parasites inside 100 WT or ire1 (-) cells. (E) WT and ire1 (-) cells were infected for 18 h and the transmigration assay was carried out for 6 h. Transmigration was determined by counting the number of infected cells normalized to noninfected cells. (±SD, n=3), **p<0.05 and ***p<0.0005. (F) WT-DCs were infected for 18 h and the transmigration assay was carried out in presence of 0.6 μM xestospongin C (XeC) for 6 h (±SD, n=3), ***p<0.001. (G-H) J774.1 macrophages were transfected with sgRNA-2 and the depletion of IRE1 was assayed by RT- qPCR and immunoblot as described in (C). (I) At 18 hpi, WT or ire1 (-) J774.1 macrophages were assayed for transmigration as described above. (±SD, n=3), **p<0.05.

### IRE1 facilitates dissemination of *Toxoplasma in vivo*

To determine the importance of IRE1 in the migration of infected DCs *in vivo*, we inoculated C57BL/6 mice intraperitoneally (i.p.) with infected WT or infected ire1 (-) DCs and measured parasite burden in the spleen by PCR at the designated time intervals over 3 days. Depletion of IRE1 in the DCs by CRISPR/Cas9 was confirmed by RT-qPCR and immunoblot analyses **(Fig. 5A and B)**. Moreover, we ascertained that there was no significant difference in parasite infection of WT and IRE1-depleted DCs **(Fig. 5C)**. *Toxoplasma* was first detected in the spleens of mice 12 h following i.p. inoculation with infected WT DCs, increasing at each time point over the 3-day period **(Fig. 5D)**. In striking contrast, appreciable levels of *Toxoplasma* dissemination of infected IRE1-depleted DCs to the spleen were not detected until 3 days following inoculation of the mice **(Fig. 5D)**. Even at the 3-day time point, the loss of IRE1 from DCs produced lowered levels of parasitemia in the spleen that were similar to those measured at 12 h of infected WT DCs. We also measured the *Toxoplasma* dissemination to the brain at 3 days, finding 200-fold fewer parasites when infected IRE1-depleted DCs were inoculated into the mice **(Fig. 5E)**. Mice inoculated with infected DCs lacking IRE1 survived significantly longer than mice receiving infected WT DCs **(Fig. 5F)**. These results demonstrate a novel role for host IRE1 in parasite pathogenesis, as IRE1 is crucial for the migration of immune cells being co-opted as “Trojan Horses” for parasite dissemination.

**Figure. 5.**
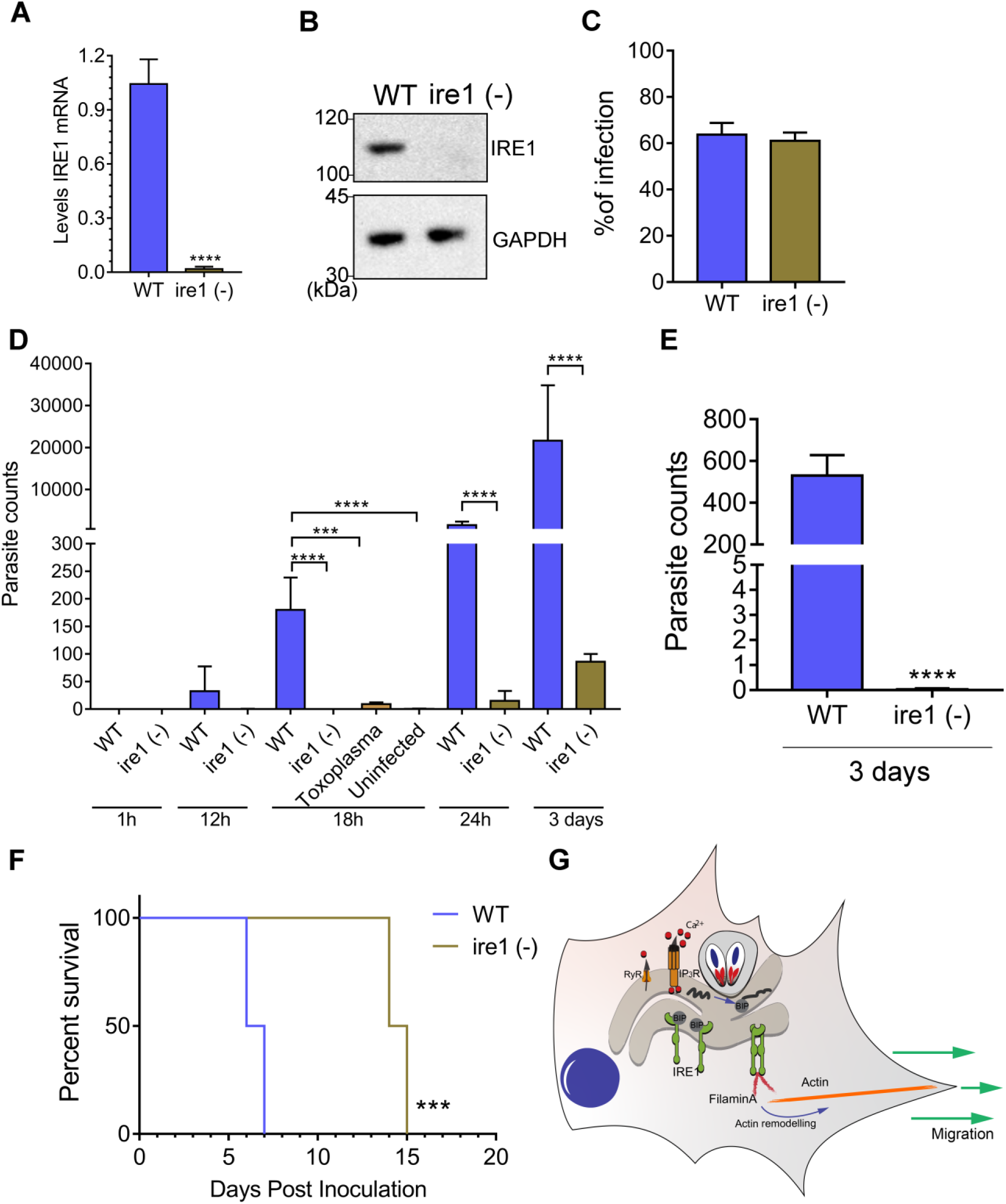
IRE1 facilitates migration of infected DCs *in vivo*. (A-B) IRE1 was depleted in bone marrow-derived DCs by CRISPR/Cas9 and loss of *IRE1* expression was assayed by RT- qPCR and immunoblot analyses. (C) WT or ire1 (-) DCs were infected for 18 h and the percentage of infection was determined by counting the number of parasites in 100 cells. (D) Infected WT and ire1 (-) cells were inoculated into mice by i.p. injection (10^6^ infected cells). At the indicated hpi, the spleen of each mouse was harvested, and the number of parasites was determined by PCR. ***p<0.005 and ****p<0.0001. (E) At 3 days post inoculation, the number of parasites was determined in the brain using PCR. ****p<0.0001. (F) Survival of C57BL/6 mice challenged with 10^6^ infected WT or ire 1 (-) DCs. ***p<0.0005. Statistical analyses by Gehan-Breslow-Wilcoxon test. (G) Model for IRE1-direct hypermigration of host cells infected by *Toxoplasma*. During infection, *Toxoplasma* triggers calcium release from the host ER, which creates ER stress and induction of the UPR, which results in enhanced IRE1 association with filamin A. Consequently, the IRE1-filamin A interaction promotes actin remodeling and host cell migration.

## Discussion

Obligate intracellular pathogens create a niche inside of their host cell that allows for parasite protection, nutrient acquisition, and the controlled release of pathogen effectors that promote infection and dissemination. *Toxoplasma* tachyzoites reside inside of a nonfusogenic parasitophorous vacuole (PV) that forms intimate contacts with host organelles and vesicles, including the ER and mitochondria [4, 5]. It is suggested that the recruitment of host organelles to the PV allows *Toxoplasma* to control critical host cell operations, including antigen presentation, nutrient production, and suppression of apoptosis [5]. In this study, we addressed consequences of the *Toxoplasma*-ER engagement on parasite infection and dissemination in the host. As illustrated in a model presented in **Fig. 5G**, we showed that *Toxoplasma* infection activates each of the UPR sensor proteins, including IRE1, via ER stress that results at least in part from release of calcium from the organelle, primarily through IP_3_R. In addition to its role in the UPR, IRE1 has recently been shown to have noncanonical functions associated with the remodeling of the cytoskeleton through direct interactions with the actin crosslinking factor filamin A [12]. We showed that *Toxoplasma* alters the morphology of its host cells through IRE1-filamin A interactions, which directs cytoskeletal remodeling that contributes to a hypermigratory phenotype that facilitated dissemination of the parasite into multiple organs of the infection host. The role of IRE1 in *Toxoplasma*-induced hypermigration is not reliant on its protein kinase or endoribonuclease activities that are central for classical UPR signaling; rather, it is IRE1 oligomerization and the C-terminal residues required for filamin A association that prove to be important.

After oral infection, *Toxoplasma* rapidly spreads from the lamina propria to distal organs using host immune cells as a vehicle for dissemination [22]. Given our discovery that IRE1 is mobilized by *Toxoplasma* to enhance hypermigratory behavior in host cells, we tested whether IRE1 is crucial to *in vivo* dissemination in a mouse model of infection. We found that depletion of IRE1 in immune cells sharply decreases the number of parasites in the spleen or brain of infected mice **(Fig. 5E)**. These results reveal a number of potential new targets for drug development aimed at thwarted spread of infection in the body.

It is noteworthy that other mechanisms have been suggested to contribute to the cell hypermotility upon *Toxoplasma* infection. For example, parasite effector protein TgWIP, TIMP- 1, and GABAergic signaling were reported to signal hypermigration of certain infected host cells [23-26]. A key question for future studies is how these different mechanisms are coordinated to induce hypermigratory activity in *Toxoplasma*-infected host cells.

## Methods

### Host cell and parasite culture

MEF (mouse embryonic fibroblast) cells were cultured in Dulbecco’s modification of Eagle’s medium (DMEM) supplemented with 10% heat-inactivated fetal bovine serum (FBS) (Gibco/Invitrogen) and penicillin/streptomycin at 37°C with 5% CO_2_. *PERK*^-/-^ cells were previously reported [27] and IRE1-deficient MEF cells were engineered by CRISPR as described below. Host cells were seeded at a density of 2×10^5^ cells/well in a 6 well plate and cultured for 18 h. Infection was performed using multiplicity of infection (MOI) of 3 with Type I or II (RH or ME49) strain *Toxoplasma* parasites, as indicated, for 18 h. The infected cells were cultured in DMEM medium supplemented with 10% heat-inactivated fetal bovine serum (FBS) (Gibco/Invitrogen) and penicillin/streptomycin at 37°C with 5% CO_2_. Cultures of DCs and J774.1 macrophages were cultivated in Roswell Park Memorial Institute (RPMI) supplemented with 10% heat-inactivated fetal bovine serum (FBS) (Gibco/Invitrogen) and penicillin/streptomycin at 37°C with 5% CO_2_ and infected as described above.

### Generation of *IRE1* knockout cells

Disruption of the *IRE1* gene in MEF cells was carried out using the CRISPR/Cas9 method [28]. Two distinct sgRNAs designed using DESKGEN™ toll (g1-TGGACACGGAGCTGACT and g2-ACACGGAGCTGACTGGG) were examined individually. The sgRNAs were prepared using the EnGen™ sgRNA Synthesis Kit (New England BioLabs), along with a sg control (g-control- CATCCTCGGCACCGTCACCC). The sgRNAs were then associated with EnGen® Spy Cas9 NLS protein (New England BioLabs) at room temperature for 20 min. MEF cells were then transfected with the bound sgRNA/Cas9 protein using the Lipofectamine CRISPRMAX Cas9 Transfection Reagent (Thermo-Fisher Scientific). After culturing the transfected cells for 48 h, 100 cells were plated in 10 mm tissue-culture dishes for cloning. Cloned *IRE1*^-/-^ MEF cells were validated by RT-qPCR using specific primers and by immunoblot using IRE1-specific antibody (Abcam-ab37073). *IRE1*^-/-^ cells were complemented with pcDNA3-derived vectors containing WT or the indicated mutant versions of *IRE1*. Briefly, the mouse cDNA sequence of *IRE1* from MEF cells was inserted into pcDNA3- EGFP plasmid (Addgene #13031), resulting in fusion proteins with the EGFP fused to the carboxy terminus of IRE1. Mutations in *IRE1* include changes inactivating the critical functions, including *kD* (kinase domain- K599A), *eD* (endoribonuclease domain- P830L), *oD* (oligomerization domain- D123P) or *c-terminal* (truncated c-terminal Δ965), were carried out using specific primers **(Supplemental Table 1)** and the Q5 Site-Directed Mutagenesis Kit (New England Biolabs). After sequence verification, the plasmids were transfected into *IRE1*^-/-^ cells using FuGENE 6 Transfection Reagent. Rescued WT and mutant IRE1 protein expression were confirmed by immunoblot and immunofluorescence microscopy **(Fig. S4A and B)**.

To generate bone marrow-derived dendritic cells (DCs), 10 × 10^6^ bone marrow cells were isolated and cultured in 6-well plate in 3 ml of complete medium (RPMI 1640 medium supplemented with 10% fetal bovine serum, penicillin, streptomycin, glutamine, 2- mercaptoethanol, 20 ng/ml granulocytes-macrophage colony-stimulating factor (GM-CSF) and 5ng/ml Interleukin 4 (IL-4) (both from Peprotech)) for 7 days. Half of the medium was replaced every two days with medium supplemented with GM-CSF and IL-4, as previously described [29]. DCs or J774.1 macrophages were transfected with IRE1-sgRNA-1 or 2, associated with EnGen™ Spy Cas9 NLS protein (New England BioLabs), using the 4D-Nucleofector™ System (Lonza) in combination with the P3 Primary Cell 4D-Nucleofector™ X Kit. After 48 h, the *IRE1* mRNA and protein levels were measured by RT-qPCR and immunoblot, respectively. Viability of DCs was examined by trypan blue staining.

### Measurement of mRNA levels

Cells were first infected with *Toxoplasma* for 2 h, washed with phosphate-buffered saline (PBS), and then cultured in DMEM at the indicated time points. RNA was isolated from cells using TRIzol LS Reagent (Invitrogen™), the cDNA was then generated using Omniscript (Qiagen) and RT-qPCR was performed using SYBR® Green Real-Time PCR Master Mixes (Invitrogen™) and the StepOnePlus Real System (Applied Biosystems™). Oligonucleotide primers used to measure each target mRNA is listed in the supplementary Table 1. Relative levels of target mRNAs from the uninfected samples were adjusted to 1 and served as the basal control value. Values of each time point were normalized to mock infection. Each experiment was performed three times, each with three technical replicates.

### Immunoblot analyses

Cells were infected with *Toxoplasma* for 2 h, washed with PBS, then cultured in DMEM for the indicated time points. The infected cells were harvested in RIPA buffer solution supplemented with cOmplete™ and EDTA-free Protease Inhibitor Cocktail (Roche). Protein quantification was performed using the Bradford Reagent (Sigma-Aldrich). Equal amounts of protein lysates were separated by SDS-PAGE and proteins were transferred to nitrocellulose filters. Immunoblot analyses were using primary antibodies- IRE1 (Abcam- ab37073), XBP1s (Cell Signaling #D2C1F), ATF6 [30], GAPDH (Abcam-ab9485), PERK (Cell Signaling #3192), followed by Amersham ECL HRP-Conjugated Antibodies secondary antibody. These antibodies and additional reagents used in the study are listed in the supplementary Table 2. Proteins were visualized in the immunoblots were visualized using FluorChem M- Multiplex fluorescence (Protein Simple). Immunoblot analyses were carried out for three independent experiments.

### Calcium measurement assay

*Toxoplasma*-infected MEF cells were washed twice with buffer A solution supplemented with glucose (120 mM NaCl, 20 mM HEPES (pH 7.4), 4.7 mM KCl, 1.2 mM NaH_2_PO_4_, 1.2 mM MgSO_4_, 1.2 mM CaCl_2_ and 10 mM glucose) [31], and then a final concentration of 5 µM of Fluo-4, AM (Thermo Fisher Scientific, F14201) was added for 15 min at 37°C. Prior to the calcium measurements, cells were washed once with buffer A solution supplemented with glucose. A Synergy (BioTek) plate reader was used to monitor the Fluo-4 AM fluorescence at 488-nm excitation and 524-nm emission wavelengths. Values derived from infected cells (*ΔF*) were divided by the resting intracellular calcium (Fo), ΔF/Fo, and the values of each time point were normalized to mock-infected cells. In parallel, live infected cells were imaged by microscopy at the same exposure and a heat map was generated using ImageJ software. To determine the activity of RyR and IP_3_R receptors, infected cells were incubated with 5 µM Mag-Fluo-4, a low-affinity Ca^2+^ indicator, then permeabilized with 10 µg/ml of saponin followed by incubation with 1.5 mM ATP to maintain Mag-Fluo-4 in the ER [19] **(Fig. S2B)**. Infected cells were treated with the indicated concentrations of caffeine (RyR, 0-200 mM) or IP_3_ (IP_3_R, 0-3 µM) (Sigma-Aldrich). A Synergy (BioTek) plate reader was used to monitor the Mag-Fluo-4 fluorescence at 490-nm excitation and 525-nm emission wavelengths [19]. Values were normalized to mock-infected cells.

### Immunofluorescence assay

Cultured cells were infected with *Toxoplasma* for 18 h, then fixed with 2.5% paraformaldehyde for 20 min and blocked with PBS supplemented with 2% BSA. Cells were permeabilized in blocking solution containing 0.01% Triton X-100 for 30 min, and incubated with primary antibody (SAG1-p30, Invitrogen) for 1 h. Secondary goat anti-rabbit Alexa-fluor 488 (Invitrogen) was applied for 1 h in the presence of Rhodamine Phalloidin (Thermo Fisher Scientific) followed by Prolong Gold antifade reagent (Invitrogen). DAPI was used to visualize host cells and parasite nuclei (Vector Labs). Images were acquired with a Leica inverted DMI6000B microscope with 63x oil immersion objective and analyzed in ImageJ. Alternatively, *IRE1*^-/-^ cells were transfected with pcDNA3 encoding IRE1 fused with EGFP at the carboxy terminus and infected with *Toxoplasma* for 18h (MOI: 3); the cells were then fixed and imaged as described above.

### Immunoprecipitation assay

The mouse cDNA sequence of filamin A from MEF cells was amplified and cloned into pcDNA3-myc plasmid (Addgene). The resulting plasmid pcDNA3- myc-FLNA was transiently transfected in the MEF cells and then the transfected cells were infected with *Toxoplasma* for 18 h (MOI: 3). Cell lysates were prepared using IP-lysis solution (0.5% NP-40, 250 mM NaCl, 30 mM Tris, 0.5% glycerol, pH 7.4, 250 mM phenylmethylsulfonyl fluoride (PMSF) supplemented with cOmplete™ and EDTA-free Protease Inhibitor Cocktail (Roche)). To immunoprecipitate myc-tagged filamin A (myc-filamin A), equal amounts of protein lysates were incubated with IgG Magnetic beads (Pierce) for 2 h, then mixed with anti-myc Magnetic beads (Pierce) overnight at 4 °C with rotation. Proteins bound to the beads were subsequently washed four times with IP-lysis solution at 4°C and then once with IP-lysis solution supplemented with 500 mM NaCl. Protein complexes were eluted at 95°C for 5 min in loading buffer solution and then separated by SDS-PAGE, followed by immunoblot analyses using specific antibodies to IRE1 (Abcam-ab37073) or Myc (Cell Signaling #2276).

### Cell migration assay

Cells were infected with *Toxoplasma* MOI 3 for 18 h and then trypsinized and counted using a hemocytometer; 2×10^4^ cells were resuspended in serum-free medium and applied to the top of a membrane coated with collagen I (rat-tail) (Gibco- A1048301). Transmigration assays were carried out using a Corning Transwell Costar apparatus (6.5 mm diameter and 8 µm pore size) as described [32]. After 18 h for MEF and 6 h for DCs and macrophages, the medium was removed, and the cells were fixed with 2.5% paraformaldehyde for 20 min. The facilitate counting of migrated cells, cells that did not migrate and remained on the upper side of membrane (unmigrated cells) were removed with a swab. The membrane was incubated with Prolong Gold antifade reagent with DAPI. Cells were counted using a Leica inverted DMI6000B microscope with 63x oil immersion objective. The transmigration was determined by numbers of migrated infected cells in 5 fields normalized to number of uninfected cells. Each transmigration assay was carried out in technical triplicate in n=3.

### *In vivo* migration assay

DCs were plated in 6-well plates (1×10^6^ cells/well) and allowed to adhere overnight. DCs were infected with *Toxoplasma* for 1 h (MOI: 3) and then washed with RPMI to remove extracellular parasites. After 18 h, infected DCs were incubated with CellTracker™ Orange CMTMR (Thermo Fisher) as described [33]. Infected DCs were intraperitoneally inoculated into 6-weeks-old female C57BL/6J mice whose spleens and brains were subsequently harvested at the indicated time points and the DC migration to the spleen was measured by the fluorescence intensity at the indicated time points using a Synergy (BioTek) plate reader at ex/em 541/565 nm **(Fig. S5C)**. Also, DNA was isolated from the spleen and brain using TRIzol (Thermo Fisher), and the number of parasites was determined by using a PCR-based method measuring levels of the parasite-specific gene region B1 as previously described [34]. After 3 days, the mice were observed twice a day and percent survival was recorded at each time point. The mice experiment, including parasite measurement by B1 were performed on blinded. The mice used in this study were housed in American Association for Accreditation of Laboratory Animal Care (AAALAC)-approved facilities at the Indiana University School of Medicine Laboratory Animal Research Center (LARC). The Institutional Animal Care and Use Committee (IACUC) at Indiana University School of Medicine approved the use of all animals and procedures (IACUC protocol number 11376).

### Quantification and statistical analysis

Quantitative data were presented as the mean and standard deviation and were derived from three biological replicates. Statistical significance was determined using One-way ANOVA with Tukey’s post hoc test and multiple t-test two-tailed using Graph Prism software. The number of biological replicates (n) and p-values are indicated in figure legends. For immunoblot analyses, the reported images are representative of at least three independent experiments. The mice survival curve was analyzed by Gehan-Breslow-Wilcoxon test for *in vivo* analysis.

## Acknowledgments

This research was supported by a research grant from National Institutes of Health (AI124723 to W.J.S. and R.C.W.). R.C.W. receives financial support from HiberCell. M.H.K. and N.S.A. were supported by funds from the Brown Center for Immunotherapy. J.M. was supported by PHS grant T32 AI060519 and the Joseph and Lucille Madri Family Scholarship. The authors would like to thank Drs. Tatiana Clemente and Stacey Gilk, as well as our lab members for helpful discussions.

## Contributions

Study design and planning: L.A., J.M., P.H.A., M.H.K, R.C.W. and W.J.S. Performed experiments and generated reagents: L.A., J.M., P.H.A. and N.S.A. Data analysis: L.A., J.M. and P.H.A. Manuscript writing: L.A., R.C.W. and W.J.S. Manuscript was drafted with input from all authors.

## Competing interests

The authors declare no competing interests.

## Supplemental Figure Legends

**Supplementary Figure. 1.**
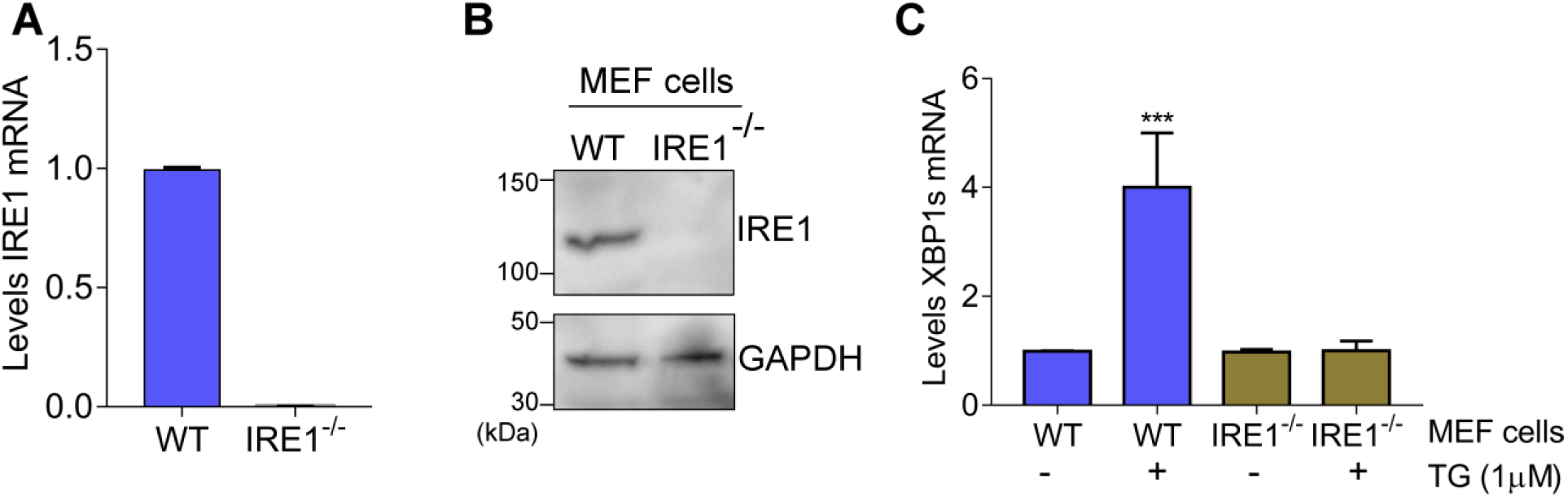
MEF cells were transfected with Cas9 bound to sgRNA targeted to IRE1 as described in the methods section; lowered *IRE1* expression resulting from the CRISPR/Cas9 gene editing was evaluated by (A) RT-qPCR and (B) immunoblot analyses. GAPDH was included as a loading control for the immunoblot analyses. (C) WT and *IRE1*^-/-^ MEF cells were treated with 1 µM thapsigargin (TG), or no stress agent, for 6 h. Cells were then harvested and the *XBP1s* mRNA levels were measured by RT-qPCR. The values of *XBP1s* mRNAs were normalized to values of total *XBP1* transcripts. (±SD, n=3) ***p<0.0005.

**Supplementary Figure. 2.**
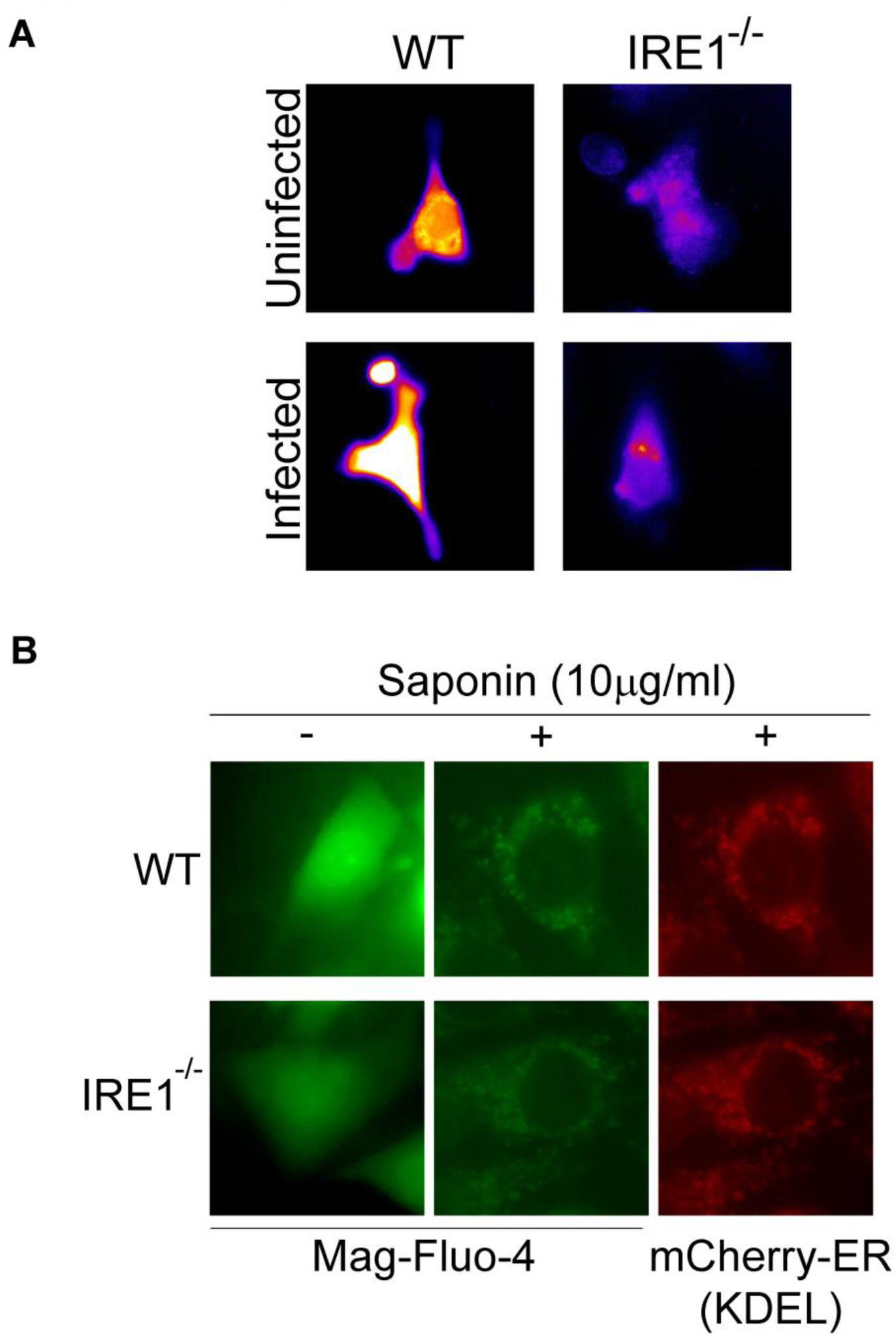
(A) MEF cells were infected with *Toxoplasma* for 18 h and then were incubated with calcium indicator Fluo-4. Fluo-4 intensity is shown as a heat map, with yellow showing the highest Fluo-4 intensity and blue showing the lowest Fluo-4 intensity. (B) WT and *IRE1*^-/-^ cells were transfected with mCherry-ER-KDEL (a marker for the ER) and infected for 18 h with *Toxoplasma*. Cells were then loaded with the low-affinity Ca^2+^ indicator Mag-Fluo-4 AM (green) and the plasma membrane was permeabilized, resulting in Mag-Fluo-4 AM retainment in the ER. Note that the Mag-Fluo-4 AM (green) co-localizes with mCherry-ER-KDEL (red).

**Supplementary Figure. 3.**
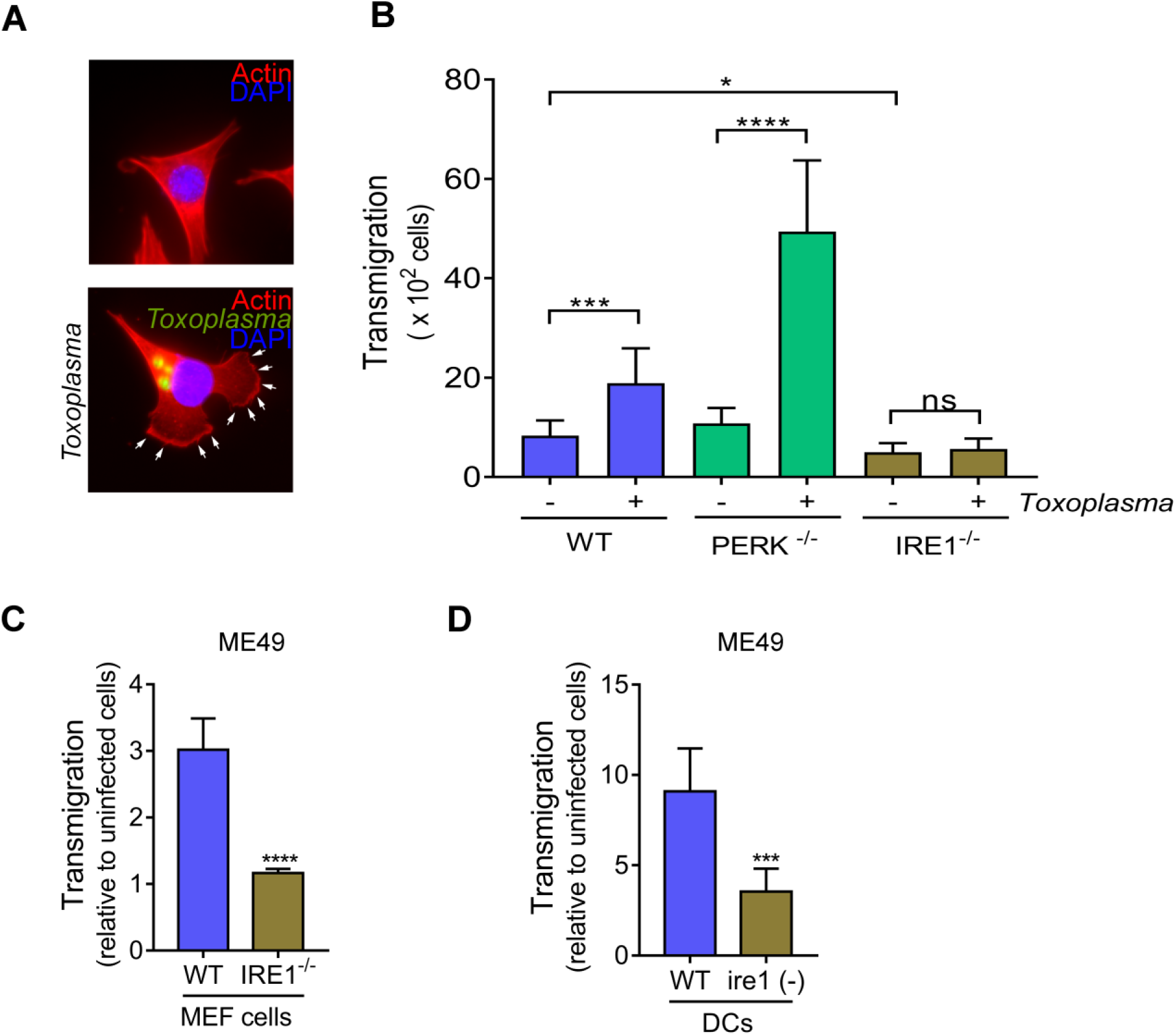
(A) WT cells were infected with *Toxoplasma* for 18 h, then fixed with paraformaldehyde and incubated with SAG1 antibody to detect parasites (green); phalloidin shows actin (red) and DAPI shows nuclei (blue). The arrows show lamellipodia at edge of cells. (B) At 18 hpi, infected and uninfected cells were trypsinized and counted, and the same number of cells were used in the transmigration assay. Transmigration was determined by counting the number of cells that transmigrated through membrane. (±SD, n=3), *p<0.05, ***p<0.0005 and ****p<0.0001. ns = not significant. (C) WT and *IRE1*^-/-^ MEF cells were infected using ME49 (type II) strain for 18h and transmigration was determined by counting the number of infected cells normalized to noninfected cells (±SD, n=3), ****p<0.0001. (D) WT and ire1 (-) DCs were infected with ME49 strain for 18 h and transmigration was determined by counting the number of infected cells normalized to noninfected cells for 6 h (±SD, n=3), ***p<0.0005.

**Supplementary Figure. 4.**
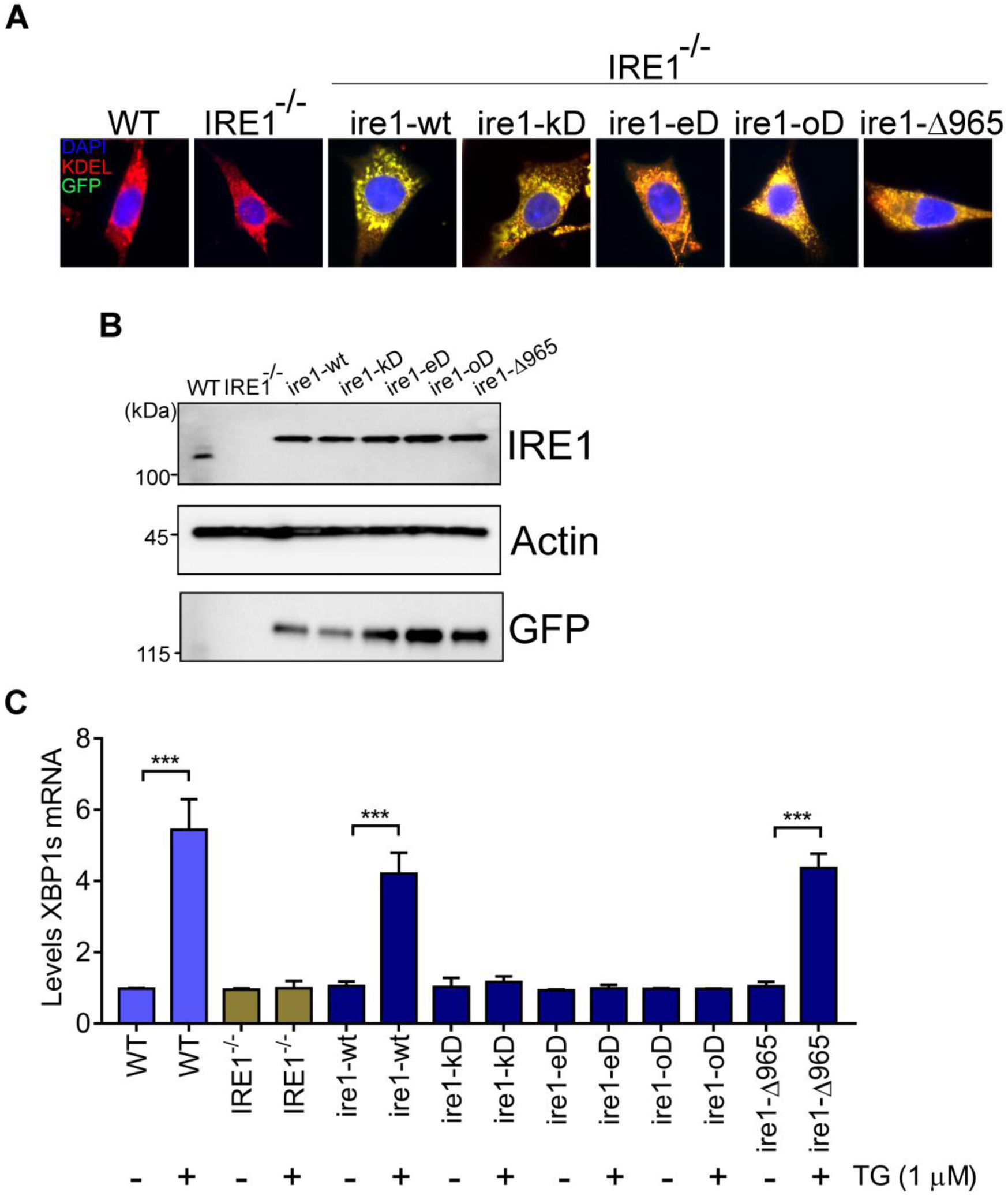
(A) *IRE1*^-/-^ MEF cells were rescued with WT and mutant versions of transient expression *IRE1* (green), then cells were fixed and stained using KDEL antibody as an ER marker (red); DAPI was used as to visualize nuclei (blue). *Ire1-wt, ire1-kD*: kinase domain dead, *ire1-eD*: endoribonuclease dead, *ire1-oD*: *ire1-Δ965* deletion of filamin A binding site [12]. (B) Lysates were prepared from the cells and IRE1, GFP, or actin protein levels were measured by immunoblot analyses using specific antibodies. (C) WT or *IRE1*^-/-^ MEF cells that were rescued with the indicated *IRE1* alleles were cultured in the presence or absence of 1 μM thapsigargin for 6 h and *XBP1s* mRNA levels were measured by RT-qPCR. Values of *XBP1s* mRNA were normalized to total *XBP1* mRNA levels for each condition. (±SD, n=3), ***p<0.0005.

**Supplementary Figure. 5.**
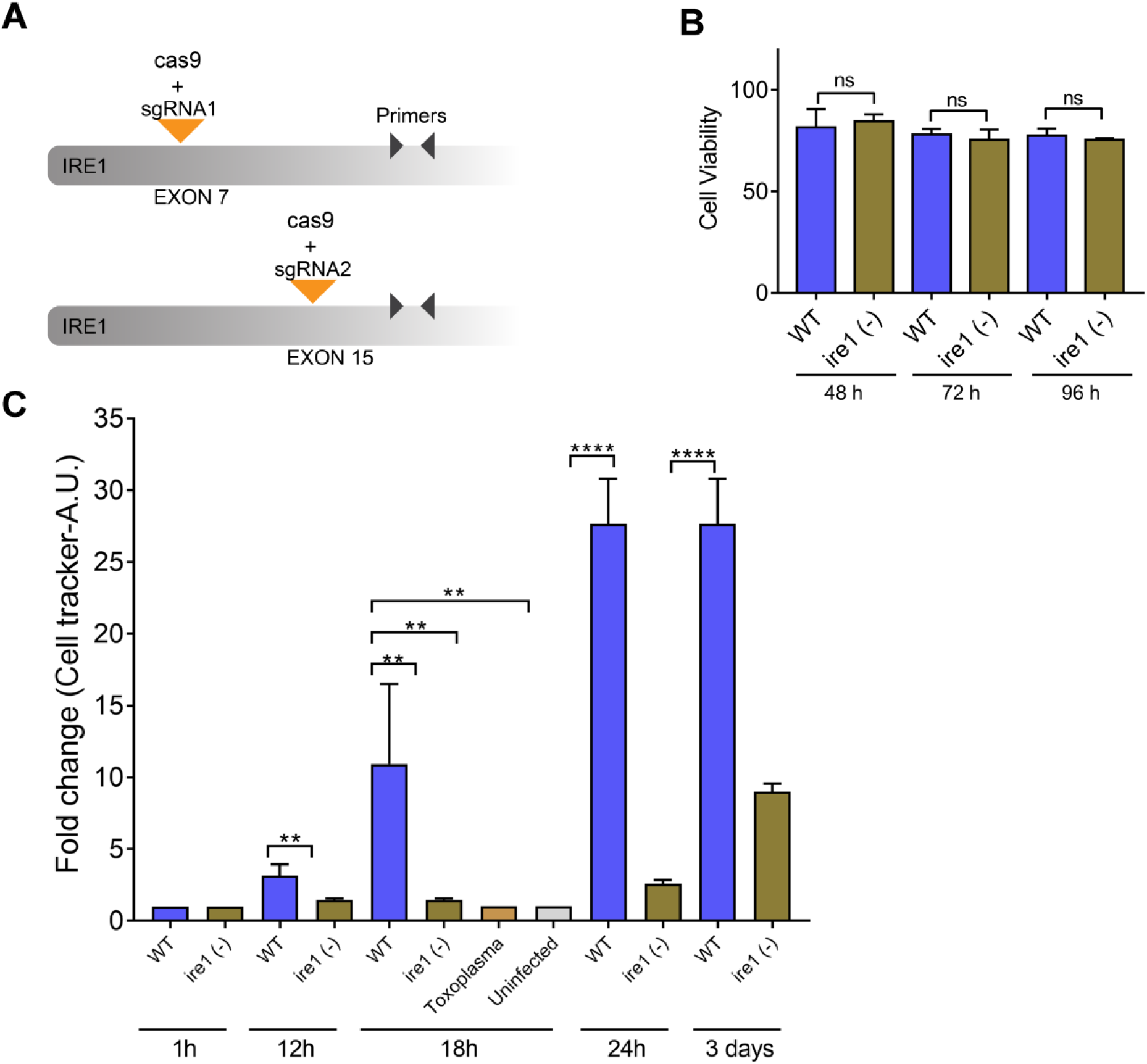
(A) Schematic of IRE1-sgRNAs (1 and 2) and the gene region that was amplified by RT-qPCR to verify loss of *IRE1* expression. (B) Viability assay of DCs at designated hours post-transfection with gRNA-IRE1 (ire1 (-)) or random gRNA (WT). (C) Infected WT and ire1 (-) DCs were incubated with cell tracker and then inoculated into mice by i.p. injection using 10^6^ infected cells. At the indicated hpi, the spleen of each mouse was harvested and the cell tracker fluorescence was measured using a plate reader. Values of fluorescence were normalized to uninfected (fold change). **p<0.01 and ****p<0.0001.

**supplementary Table 1.**
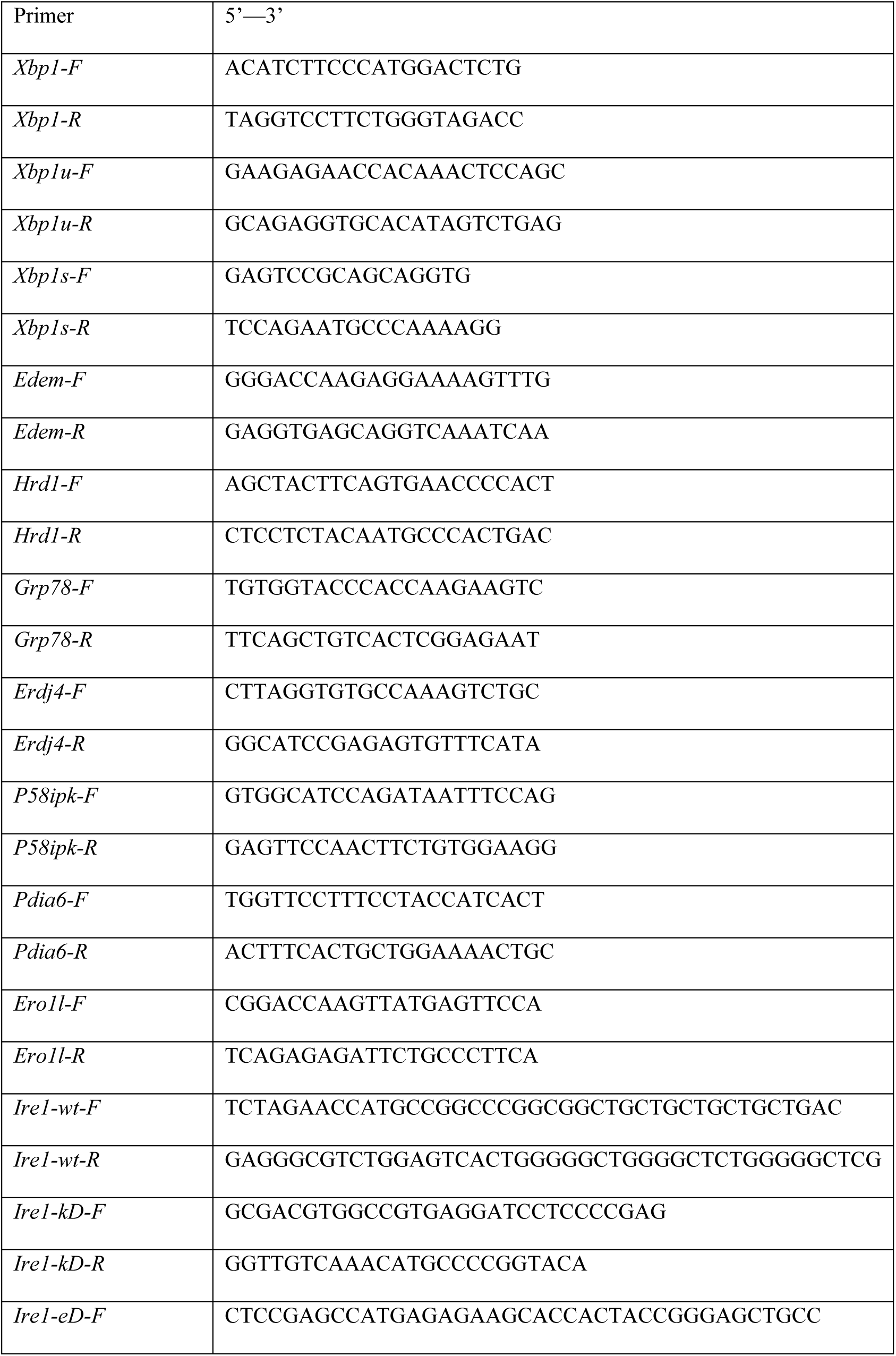
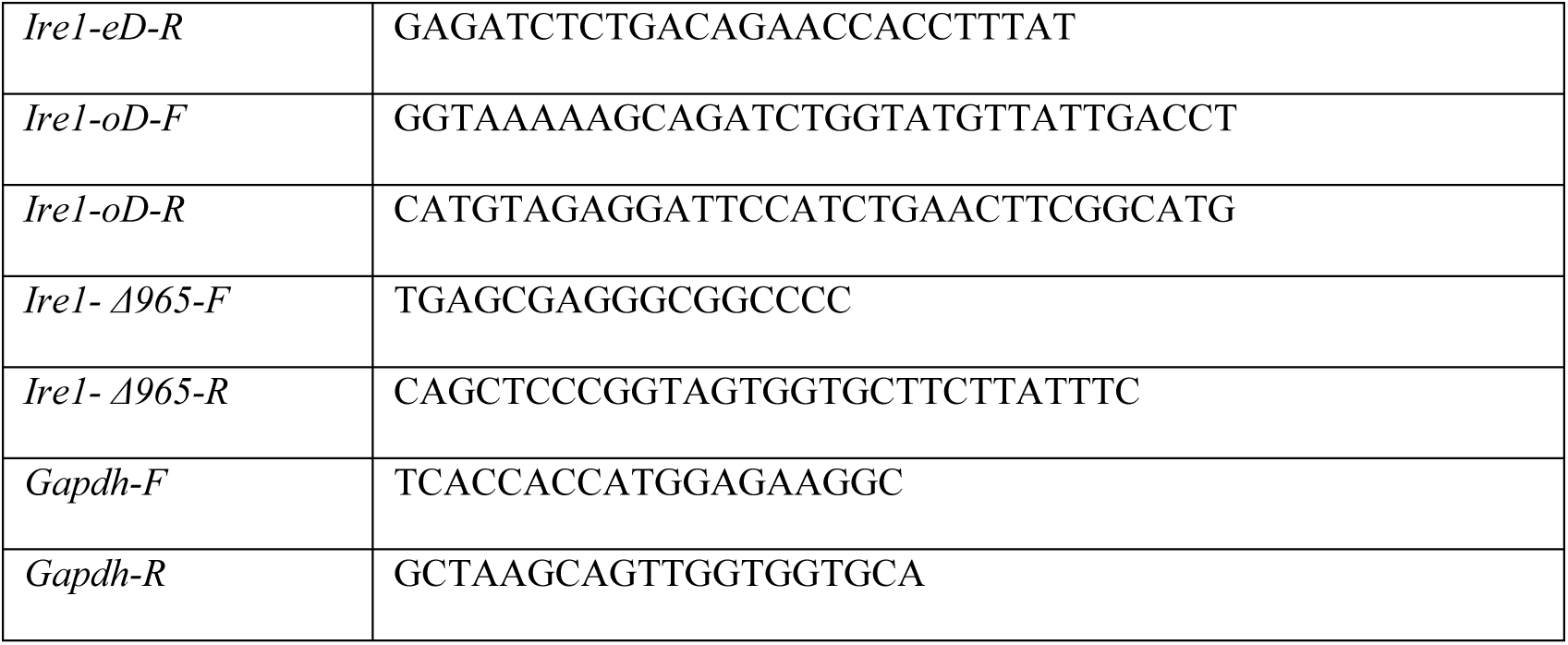
Oligonucleotide primers used in this study.

**supplementary Table 2.** Reagents used in this study.

| Reagent | Company names |
| --- | --- |
| **Antibody** |  |
| PERK-P | Phospho-PERK (Thr980) (16F8) Rabbit mAb #3179- Cell Signaling |
| PERK | PERK (C33E10) Rabbit mAb #3192- Cell Signaling |
| ATF6 | [30] |
| XBP1s | XBP-1s (D2C1F) Rabbit mAb #12782- Cell Signaling |
| GAPDH | ab9485-Abcam |
| Phalloidin | R415-Thermo Fisher Scientific |
| SAG1 | Toxoplasma gondii P30 Monoclonal Antibody (P30/3)- Thermo Fisher Scientific |
| IRE1 | ab37073-Abcam |
| Myc | Myc-Tag (71D10) Rabbit mAb #2278-Cell Signaling |
| GFP | clone GFP-20-Sigma Aldrich |
| Filamin A-P (S2152) | Phospho-Filamin A (Ser2152) Antibody #4761-Cell Signaling |
| Filamin A | Filamin A Antibody #4762-Cell Signaling |
| **Reagent** |  |
| Fluo-4, AM | F14201-Thermo Fisher Scientific |
| Ry | 1329-Tocris |
| XeC | (-)-Xestospongin C- 1280-Tocris |
| IP<sub>3</sub> | D-myo-Inositol 1,4,5-trisphosphate- 1482-Tocris |
| caffeine | C53-Sigma Sigma Aldrich |
| Mag-Fluo-4 | M14206- Thermo Fisher Scientific |
| PERKi | GSK2656157- 5.04651-Sigma Aldrich |
| 4μ8c | 4479-Tocris |
| Ceapin-A7 | SML2330-Sigma Aldrich |
| KIRA6 | 19151-Cayman Chemical |
| Collagen I | A1048301-Gibco |
| Thapsigargin | 1138-Tocris |
| DAPI | D9542-Sigma Aldrich |

